# Quantitative Assessment of Protein-DNA Interactions via SYBR Green Fluorescence

**DOI:** 10.1101/2025.06.06.658291

**Authors:** Jingxin Wang, Jingwen Wang, Zhibo Wang, Pengyu Wang, Shilin Sun, Xiaofu Li, Zhen Tian, Ruibing Xu, Yifan Shi, Yucheng Wang

## Abstract

Protein-DNA interactions are crucial for cellular processes, but current quantification methods lack sensitivity. We developed the Real-time PCR-based DNA Binding Assay (RP-DBA) to detect and quantify these interactions. The target protein, expressed as a Strep-tag II fusion, is purified and incubated with double-stranded DNA probes containing 4 bp 3’ overhangs. Protein-DNA complexes are immobilized on Strep-Tactin beads, washed, and eluted. A complementary single-stranded DNA amplification arm is added, extended by Taq polymerase, and quantified via qPCR with SYBR Green. RP-DBA enables real-time kinetic analysis and is 4– to 10-fold more sensitive than EMSA, depending on amplification arm length (30–90 bp). Its simplicity, speed, accuracy, and high-throughput potential make it a valuable tool for advancing molecular biology research.

## Introduction

Life is characterized by the relationships between molecules rather than being a property of any single molecule (Cozzolino et al, 2021). Among the interaction between moleculars, the interaction between proteins and DNA plays a crucial role in the sustainability of life and the functionality of living cells. It serves as the foundation for cellular functions, controlling processes such as replication, recombination, transcription, and repair. Certain DNA-binding proteins are also associated with specialized regions within the genome, including telomeres and centromeres, thereby participating in the maintenance of chromosome structure and the regulation of chromosome condensation and pairing (Gustafsdottir et al, 2007). Additionally, these interactions are involved in various processes such as viral infections, DNA packaging, and DNA modifications (Ferraz et al, 2021). Furthermore, the interaction between proteins and DNA is critical for gene expression regulation, including the activation and repression of genes by proteases involved in chromatin modifications. For instance, transcription factors (TFs) bind to specific cis-regulatory elements, thereby inhibiting or activating gene transcription and participating in various signaling processes related to cell differentiation, development, and environmental changes (Savinkova et al, 2021).

Given the pivotal role of protein-DNA interactions in biological processes, extensive research efforts have been dedicated to the development and optimization of diverse methodologies for their investigation. These techniques can be systematically classified into in vitro and in vivo approaches based on the experimental context. In vitro methodologies encompass a range of established techniques, including DNA footprinting, electrophoretic mobility shift assay (EMSA), surface plasmon resonance (SPR), enzyme-linked immunosorbent assay (ELISA), fluorescence resonance energy transfer (FRET), and in situ hybridization. Conversely, in vivo approaches comprise yeast one-hybrid assays, chromatin immunoprecipitation (ChIP), systematic evolution of ligands by exponential enrichment (SELEX), and reverse ChIP (Li and Herskowitz, 1993; Roulet et al, 2002; Furey, 2012; Wen et al, 2020). The cumulative advancement of these methodologies has substantially enhanced our understanding of protein-DNA interactions.

In the context of high-throughput studies, in vitro methods are generally preferred due to their capacity for systematic sample processing under controlled experimental conditions, which facilitates rapid screening and comprehensive data analysis. Among these techniques, SPR has emerged as a particularly valuable tool, offering exceptional sensitivity and the unique ability to simultaneously measure binding affinities and kinetics. However, the application of SPR is constrained by several technical limitations, including non-specific binding, stringent sample purity requirements, surface-induced conformational changes, quantitative analysis challenges, high instrumentation costs, complex data processing requirements, and substantial sample consumption. These factors collectively influence the precision and practical implementation of SPR in protein-DNA interaction studies.

As a cornerstone technique in the field, EMSA is one of the most historically significant and widely used in vitro methods for investigating protein-DNA interactions, developed by Fried and Crothers (Fried and Crothers, 1981) and Garner and Revzin (Garner and Revzin, 1981). EMSA is a reliable and straightforward method for qualitatively assessing protein-DNA binding, characterized by its simplicity, sensitivity, and powerful capabilities (Pagano et al, 2011). It can detect the binding of proteins using low concentrations of both proteins and nucleic acids. This method can analyze the binding of proteins to oligonucleotides of various sizes, with probe lengths ranging from a few dozen nucleotides to thousands of nucleotides or base pairs. Additionally, it can detect nucleic acids of different structures, including single-stranded DNA, RNA, double-stranded DNA, and even triple-stranded or quadruple-stranded nucleic acids (Ferraz et al, 2021). Despite its extensive application, EMSA is associated with several notable limitations that impact its utility in contemporary research. The technique is characterized by a time-intensive experimental protocol and substantial sample requirements, particularly in terms of high-quality protein and nucleic acid materials (Gustafsdottir et al, 2007). Furthermore, the methodological complexity of polyacrylamide gel electrophoresis and subsequent visualization procedures presents technical challenges. Perhaps most significantly, EMSA is inherently limited in its capacity to analyze multiple protein-DNA/RNA interactions simultaneously, rendering it suboptimal for high-throughput applications, especially in the context of complex biological systems or large-scale interaction networks (Gustafsdottir et al, 2007).

To address these issues, we developed the Real-time PCR-based DNA Binding Assay (RP-DBA), a rapid method for studying protein-DNA interactions. This technique involves fusing the target protein with a tagged protein and incubating it with DNA. Proteins are then captured using magnetic beads that bind to the tagged protein, allowing for the collection of both the protein and its associated DNA. After washing to remove non-specific impurities, the DNA is eluted and quantitatively analyzed using SYBR Green dye. This method eliminates the need for labeling binding partners and avoids polyacrylamide gel electrophoresis, membrane transfer, or immunostaining. It enables quantitative detection of protein-DNA binding affinity and offers a robust platform for research. Importantly, it has significant potential for high-throughput applications, particularly when integrated with automation or advanced detection technologies.

## Materials and Methods

### Prokaryotic Protein Expression and Purification

We selected proteins from diverse species, including birch (*Betula platyphylla*) and *Arabidopsis thaliana*, for prokaryotic expression. The coding sequences (CDSs) of these proteins were fused with the strep tagII sequence and cloned into the pMAL-C5x plasmid encoding maltose-binding protein (MBP), generating recombinant pMAL-C5x-protein vectors. These constructs were transformed into *E. coli* strain 2523. Primers used are listed in Supplementary Table S1.

Transformed *E. coli* colonies were inoculated in LB liquid medium supplemented with ampicillin (50 mg/L) and cultured at 37°C until the optical density at 600 nm (OD_600_) reached 0.5–0.6. Protein expression was induced by adding 0.3 mM isopropyl β-D-1-thiogalactopyranoside (IPTG), followed by continued incubation for 3 hours. Cells were harvested by centrifugation at 3,000 × *g* and resuspended in chemotaxis buffer (20 mM Tris-HCl, pH 7.4, 0.2 M NaCl, 1 mM EDTA, 1 mM dithiothreitol DTT).

Resuspended cells were lysed using a sonicator (Scientz, Ningbo, China) at 180 W power output with a cycle of 5 seconds of sonication followed by 10 seconds of rest, repeated for 10 minutes. The lysate was centrifuged at 12,000 × *g* for 10 minutes, and the supernatant was collected. A portion of the supernatant was analyzed by sodium dodecyl sulfate-polyacrylamide gel electrophoresis (SDS-PAGE). Proteins were affinity-purified using amylose resin. After washing the resin, bound proteins were eluted with 10 mM maltose and stored at –80°C for subsequent use.

### Probe Primer Design

Probes were designed based on the DNA motif sequences bound by transcription factors. Each probe contains three tandem repeats of the DNA motif sequence, separated by spacer sequences. Single-stranded probes, after annealing, form sticky ends with NNNN sequences at both 3’-overhangs. These sticky ends are designed to hybridize with a complementary single-stranded DNA fragment (referred to as the signal amplification arm) in the quantitative PCR (qPCR) system. During PCR extension, the hybridization between the probe and the amplification arm generates a longer double-stranded DNA product, thereby enhancing the SYBR Green fluorescence signal intensity. Competitive probes share identical sequences with the normal probes but lack the sticky ends, preventing hybridization with the signal amplification arm. Consequently, increasing concentrations of competitive probes progressively reduce fluorescence intensity by competing for binding to the target protein. All probe types (normal, competitive, and negative controls) share the same sticky-end sequences, enabling amplification with a universal complementary DNA arm (see Supplementary Table S2 for synthesis details). Negative control probes (containing non-binding DNA sequences) retain the 3’-overhangs with sticky ends identical to normal probes, allowing PCR amplification with the signal amplification arm to establish baseline fluorescence (probe design schematics are shown in Figure 1).

**Figure 1:**
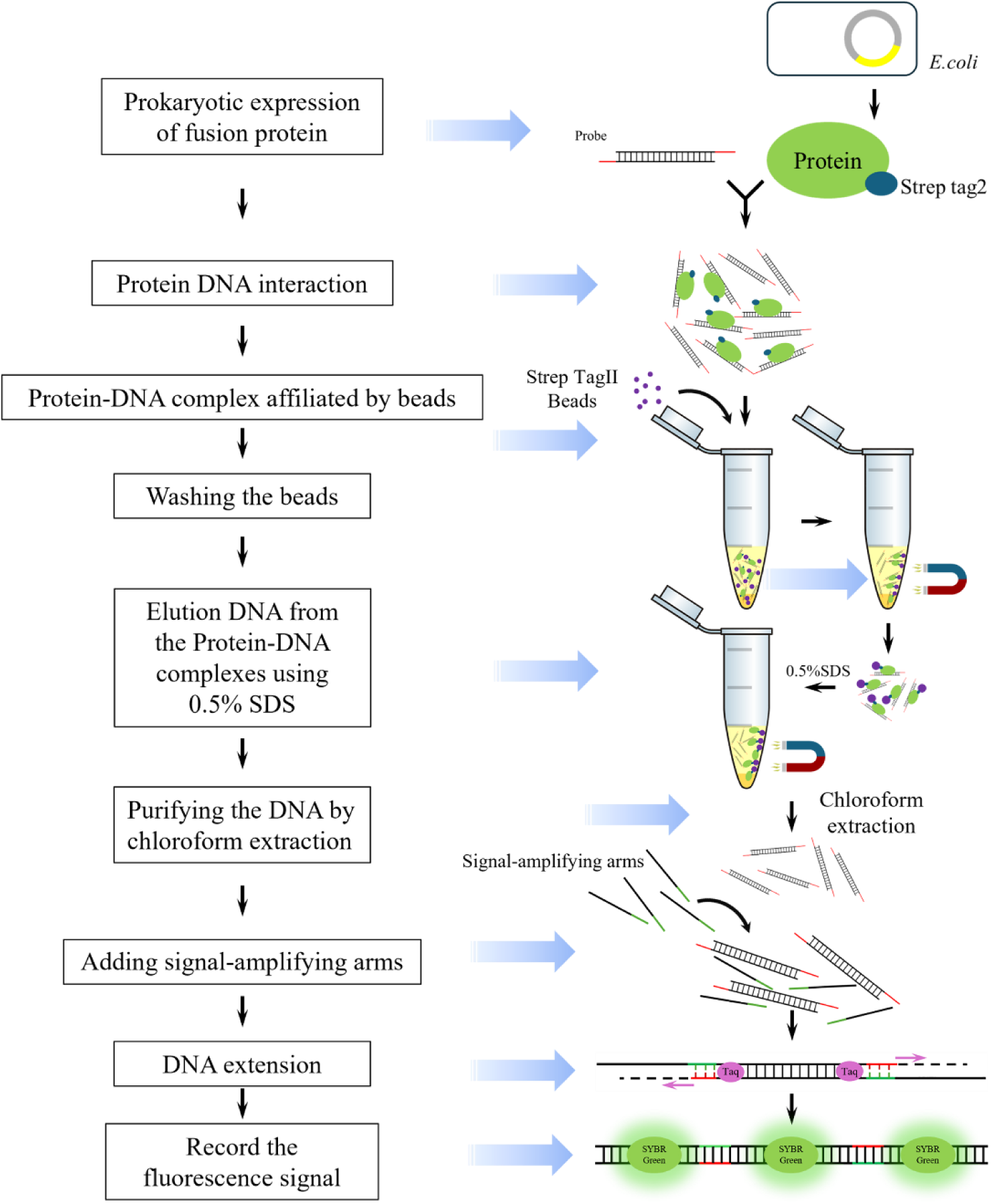
Principle and Workflow of Real-time PCR-based DNA Binding Assay (RP-DBA) The RP-DBA method involves the synthesis of probes, where the DNA sequences that bind to the protein can be repeated 2-3 times or more, typically with a length of approximately 20-30 base pairs, and feature a 3’ overhanging sticky end. The target protein is expressed in prokaryotic systems, and the protein is fused with tags such as Strep-tag II. The target protein is then incubated with the probes. After incubation, the protein is purified using affinity magnetic beads, such as Strep-Tactin beads, which simultaneously isolate the DNA that is bound to the protein. Following the washing of the beads, DNA is eluted using a 0.5% SDS solution, and chloroform extraction is performed to remove SDS for DNA purification. Quantitative PCR reagents and single-stranded DNA primers (signal amplification arms) are added to the DNA solution, which can hybridize with the sticky ends of the probes. A round of PCR extension is then conducted, using the signal amplification arms as templates to form double-stranded DNA. Finally, real-time quantitative PCR is employed to detect the fluorescence signal intensity, enabling the quantification of DNA that is bound to the protein.

### Real-time PCR-based DNA Binding Assay (RP-DBA) Protocol

The Step-by-Step Procedure was as the follows: **(1) Protein-Probe Binding Reaction:** Purified transcription factor protein (10 μL, 60 ng/μL) was mixed with 10 μL of probe (10 μM) and 20 μL of 5× EMSA/Gel-Shift Binding Buffer (100 mM HEPES-KOH, 250 mM KCl, 10 mM MgCl_2_, 12.5 mM spermidine, 5% Ficoll 400, 0.1 mM Zn(Ac)_2_, 2.5 mM DTT, 0.5 μg/mL BSA). Ultra-pure water was added to adjust the final reaction volume to 100 μL. **(2) Magnetic Bead Preparation:** 250 μL of Strep-Tactin magnetic beads (Beaver, Suzhou, China) were washed with 250 μL of 1× EMSA/Gel-Shift Binding Buffer. Washed beads were resuspended in 250 μL of 1× EMSA/Gel-Shift Binding Buffer and divided equally into 5 aliquots (50 μL each). Each aliquot was added to the protein-probe mixture and incubated for 60 minutes at room temperature with gentle agitation. **(3) Bead Washing:** Beads were sequentially washed with the following buffers (1 mL each): ①Low Salt Wash Buffer: 20 mM Tris (pH 8.0), 2 mM EDTA, 0.1% SDS (w/v), 1% Triton X-100 (v/v), 150 mM NaCl. ②High Salt Wash Buffer: 20 mM Tris (pH 8.0), 2 mM EDTA, 0.1% SDS (w/v), 1% Triton X-100 (v/v), 500 mM NaCl. ③LiCl Wash Buffer: 20 mM Tris (pH 8.0), 1 mM EDTA, 0.5% NP-40 (v/v), 0.5% sodium deoxycholate (w/v), 250 mM LiCl. ④TE Buffer: 10 mM Tris (pH 8.0), 1 mM EDTA. **(4) Probe Elution:** DNA-protein complexes were eluted using 100 μL of TE Buffer containing 0.5% SDS. Beads were thoroughly resuspended, and the supernatant was collected using a magnetic rack. To remove SDS, the eluted probes were extracted twice with chloroform by vortexing for 10 seconds and then centrifuging at 12,000 × g for 5 minutes to collect the aqueous phase. **(5) qPCR Amplification:** ①Purified DNA (9 μL) was mixed with 10 μL of ChamQ SYBR Green qPCR Master Mix (Vazyme, Nanjing, China) and 1 μL of single-stranded DNA amplification arm (10 μM). ②The mixture was incubated at 42°C for 3 minutes to facilitate annealing, followed by fluorescence detection using a quantitative PCR instrument.

**Controls and Replicates:** ①Negative control: Non-binding DNA sequence with identical sticky ends. ②Competitive probe controls: Competitor probes (identical to normal probes but lacking sticky ends) were added at 1×, 10×, and 20× molar excess relative to the normal probe. ③All reactions were performed in triplicate technical replicates.

**Probe Sequences:** Detailed sequences for all RP-DBA probes are provided in Supplementary Table S2.

### Chemiluminescent EMSA

Probes were end-labeled with Biotin using the Biotin 3′ End DNA labeling kit (Pierce Biotechnology, Rockford, IL, USA). The reaction system consisted of 10 μL of TdT Buffer (5×), 5 μL of probe (1 μM), 5 μL of Biotin-11-dUTP (5 μM), and 1 μL of Terminal Deoxynucleotidyl Transferase (TdT) (10 U/μL), with a total reaction volume of 50 μL. The reaction was incubated at 37°C for 30 minutes. The labeled probes were then co-incubated with purified proteins to perform chemiluminescent EMSA. After probe-protein incubation, the samples were electrophoresed on a non-denaturing polyacrylamide gel (PAG) at 100 V, using 0.5× TBE as the running buffer. The resolved complexes were transferred onto a nylon membrane via electroblotting. UV crosslinking was performed for 3–10 minutes. Streptavidin-Horseradish Peroxidase (HRP) was added to bind Biotin, followed by chemiluminescent development. Signals were visualized using the Tanon 5200 Chemiluminescent Imaging System (Tanon Science and Technology, Shanghai, China). All probe sequences used in EMSA are listed in Supplementary Table S3.

### Sensitivity Study of the Protein-DNA Binding Assay (RP-DBA)

Protein samples were equally divided into three aliquots and subjected to serial dilution (1×, 0.5×, 0.2×, 0.1×). The diluted protein samples were applied to two detection methods: conventional EMSA and RP-DSA. By comparing the decay trends of signal intensity between the two methods at varying protein concentrations, the detection sensitivity of RP-DBA was systematically evaluated.

### Analysis of DNA-Protein Binding Kinetics

To quantify binding kinetic parameters, under fixed probe concentration conditions, nine time gradients were set up: 0, 5, 10, 15, 20, 25, 30, 35, 40, 45 minutes for protein-DNA complex incubation. At each time point, the reaction was terminated using magnetic bead separation, and subsequent experimental procedures followed the protocol described in Section 2.3, followed by fluorescence detection. Three replicates were performed.

### Study on Signal Sensitivity of Probes with Different Arm Lengths

This study systematically compared the signal differences between probes modified with arm extensions of varying lengths and unmodified probes. The protein-DNA binding reaction system and experimental conditions were identical, with the critical distinction as follows: after protein-DNA complex formation, 30 bp (used in this study), 60 bp, and 90 bp specific arm extensions (Supplementary Table S2) were introduced into the quantitative PCR reaction mixture for the experimental groups to evaluate their impact on enhancing signal sensitivity, while no arm was added to the control group. Subsequent procedures (e.g., separation, washing, and detection) were rigorously standardized. Quantitative analysis was performed using a real-time quantitative PCR (qPCR) fluorescence detection system. All probe sequences used in this study are listed in Supplementary Table S2.

### Histograms and statistics

The histograms were generated by using Prism8.0 (GraphPad). The statistical significance of intergroup differences was evaluated using one-way analysis of variance (ANOVA) test.

## Results

The principle of Real-time PCR-based DNA Binding Assay (RP-DBA) is as follows: we utilize the property of SYBR Green that binds to dsDNA and emits fluorescence, allowing it to be detected by quantitative PCR instruments. First, similar to methods such as yeast one-hybrid and EMSA, we synthesized DNA elements that bind to the protein. The sequence of this element is formed by the annealing of two single-stranded DNAs, which have complementary sticky ends and protruding 3’ ends on both sides. These sticky ends are used to pair with a single-stranded DNA sequence (signal amplification arm), which is then extended into double-stranded DNA in the PCR system, enhancing the detection signal (Figure 1).

Next, we express and purify the target DNA binding protein, which needs to be fused with a Strep tag II for tight affinity with magnetic beads. The purified protein is incubated with an excess of probes to allow the probes to bind to the protein. Then, magnetic beads are used to isolate the protein-DNA complexes, and non-specific binding components are removed through washing. Following this, a TE solution containing 0.5% SDS is used to incubate with the magnetic beads to release the DNA bound to the protein. At this point, the DNA needs to undergo chloroform extraction to remove the SDS. Subsequently, the obtained DNA is incubated with the quantitative PCR mix, along with the signal amplification arm (a single-stranded DNA sequence). This sequence binds to both ends of the target DNA through the complementarity of the sticky ends, allowing Taq polymerase to amplify the single-stranded DNA template to form double-stranded DNA (only one round of PCR) to increase the concentration of the fluorescent signal (Figure 1).

To set up control experiments, we designed two types of controls: the first is a competitive probe, which has the same DNA sequence as the normal probe but lacks sticky ends, thus preventing signal amplification after purification. This probe will produce a significant signal difference compared to the normal probe. The second is a negative control, which consists of a DNA sequence that does not bind to the target protein but has the same sticky ends as the probe. This negative control can also complement the signal amplification arm and undergo PCR amplification, thereby amplifying the signal to compare the specific binding of the protein to the DNA sequence.

It is important to note that the sticky ends and their paired single-stranded DNA sequences (signal amplification arm) are unrestricted, as they do not bind to the target protein (assuming we are studying double-stranded DNA-binding proteins). Additionally, to reduce costs, errors, and operational complexity, we designed the sticky end sequences on both DNA 3’ends to be consistent, allowing the use of the same signal amplification arm. The amount of protein used in each reaction should remain consistent (ensured by controlling the volume of magnetic beads), while the probes should be kept in a saturated state to improve experimental accuracy. Ultimately, the main criterion for determining specific binding is whether the fluorescent signal is significantly higher than that of the control group.

### Real-time PCR-based DNA Binding Assay (RP-DBA)

To investigate the specific binding of RP-DBA to DNA, we selected three transcription factors derived from birch or Arabidopsis and expressed them in a prokaryotic system. Subsequently, we analyzed the binding capacity of these transcription factors to their target element sequences. We initially used the RP-DBA method to evaluate their binding to the corresponding target DNA elements, utilizing an amplification arm length of 30 bp. To ensure the reliability and reproducibility of the experimental results, each experiment was conducted in triplicate. The results demonstrated that all three transcription factors exhibited specific binding to the target DNA elements, generating strong fluorescent signals in quantitative PCR detection. As the concentration of the competitive probe increased, the fluorescent signal in the experimental group gradually diminished, creating a clear competitive signal gradient (Figure 2A-F), indicating that the binding of the transcription factors to the target DNA is competitive. Furthermore, the fluorescent signal from the negative control (DNA sequences that do not bind to the target protein) was significantly lower than that of the experimental group, further excluding the possibility of non-specific binding (Figure 2A-F).

**Figure 2:**
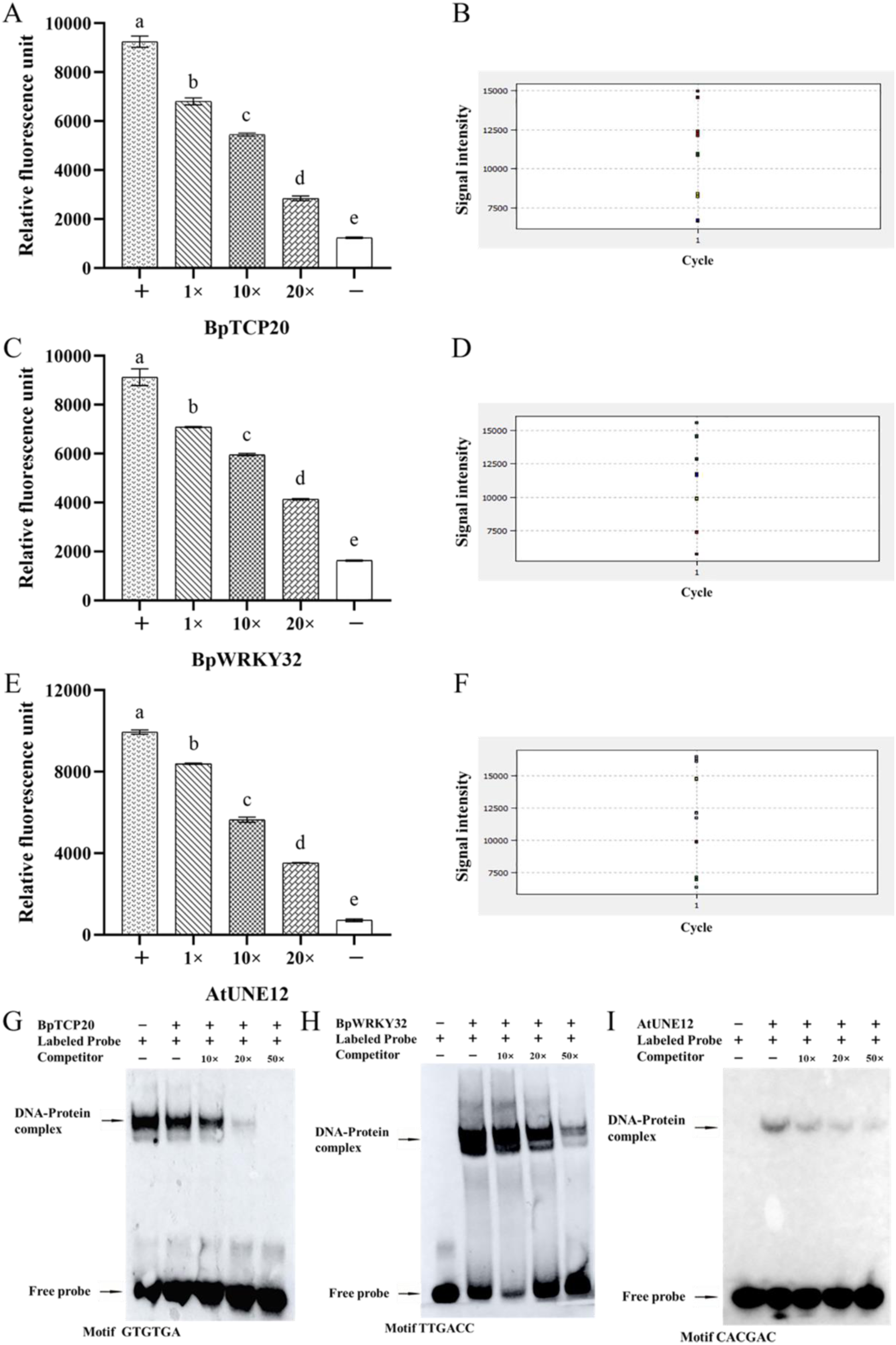
Investigation of Protein-DNA Interactions Using RP-DBA. A, C, and E: the quantity of DNA bound to the target proteins, as determined by fluorescence signal intensity. These three proteins are BpTCP20 (A) and BpWRKY2 (C) from birch, and AtUN12 (E) from Arabidopsis thaliana. B, D, and F: The original fluorescence signal intensity graphs for the DNA-protein binding, generated by the quantitative PCR instrument, corresponding to BpTCP20 (B), BpWRKY2 (D), and AtUN12 (F), respectively. The signal intensity values in panels A, C, and E represent the signal values in panels B, D, and F after subtracting the background values (using an equal volume of water as a control). The error bars indicate the standard deviations of the mean measurements (n= 3). The statistical significance of intergroup differences was evaluated using one-way analysis of variance (ANOVA) test. Letters a-e indicate significant differences (P < 0.01). An amplification arm with a length of 30 bp was used. F, G, and H: Validation of the interactions between the aforementioned proteins and DNA using EMSA, corresponding to BpTCP20, BpWRKY2, and AtUN12, respectively. The EMSA probes are end-labeled with biotin, and chemiluminescence is employed to detect the binding of proteins to DNA.

To quantify the experimental results, we calculated the signal differences among the experimental group, the competitive probe group, and the negative control group. We also used ultrapure water as a background control to minimize the influence of background fluorescence. After subtracting the background signal, we computed the signal values, average signal values, and the significance of the differences (p < 0.01) for each group. The results indicated that the fluorescent signal in the experimental group was significantly higher than that in both the competitive probe group and the negative control group (p < 0.01) (Figure 2A-F), demonstrating a high degree of specificity in the binding of the transcription factors to the target DNA. Additionally, the results from the three independent replicates were highly consistent, with small standard deviations (Figure 2A-F), indicating that the RP-DBA method possesses good reproducibility.

To further validate the reliability of the RP-DBA results, we conducted electrophoretic mobility shift assays (EMSA) as a parallel detection method for the binding of transcription factors to the target DNA. The results from RP-DBA were consistent with those obtained from EMSA (Figure 2G), providing additional evidence for the accuracy and reliability of the RP-DBA technique.

In summary, the RP-DBA method effectively detects the specific binding of transcription factors to target DNA, exhibiting high reproducibility and reliability. By incorporating competitive probes and negative controls, this method can clearly distinguish between specific and non-specific binding, offering a sensitive and efficient technological approach for studying protein-DNA interactions.

### Kinetic Study of Protein-DNA Interactions Using RP-DBA Technology

Previously, we conducted ChIP-Seq analyses of the bHLH transcription factor from birch (public database accession number: PRJNA1247694) and performed peak calling using the MACS2 software (q-value < 0.05) to identify significantly enriched regions (peaks). Subsequently, we employed the *de novo* motif prediction tool MEME Suite (v5.5.1) to mine conserved motifs within the ±100 bp sequences surrounding the peak centers. Through motif enrichment analysis using HOMER, we discovered several novel DNA elements, identifying multiple potential binding sites for bHLH. We randomly selected three of these candidate elements for further study (Figure S1). Initially, we investigated the binding of these elements to bHLH using the RP-DBA method, which revealed that all three elements bound to bHLH (Figures 3A-F). We further validated the reliability of the RP-DBA method by conducting EMSA on these three elements (Figure 3G-I).

**Figure 3:**
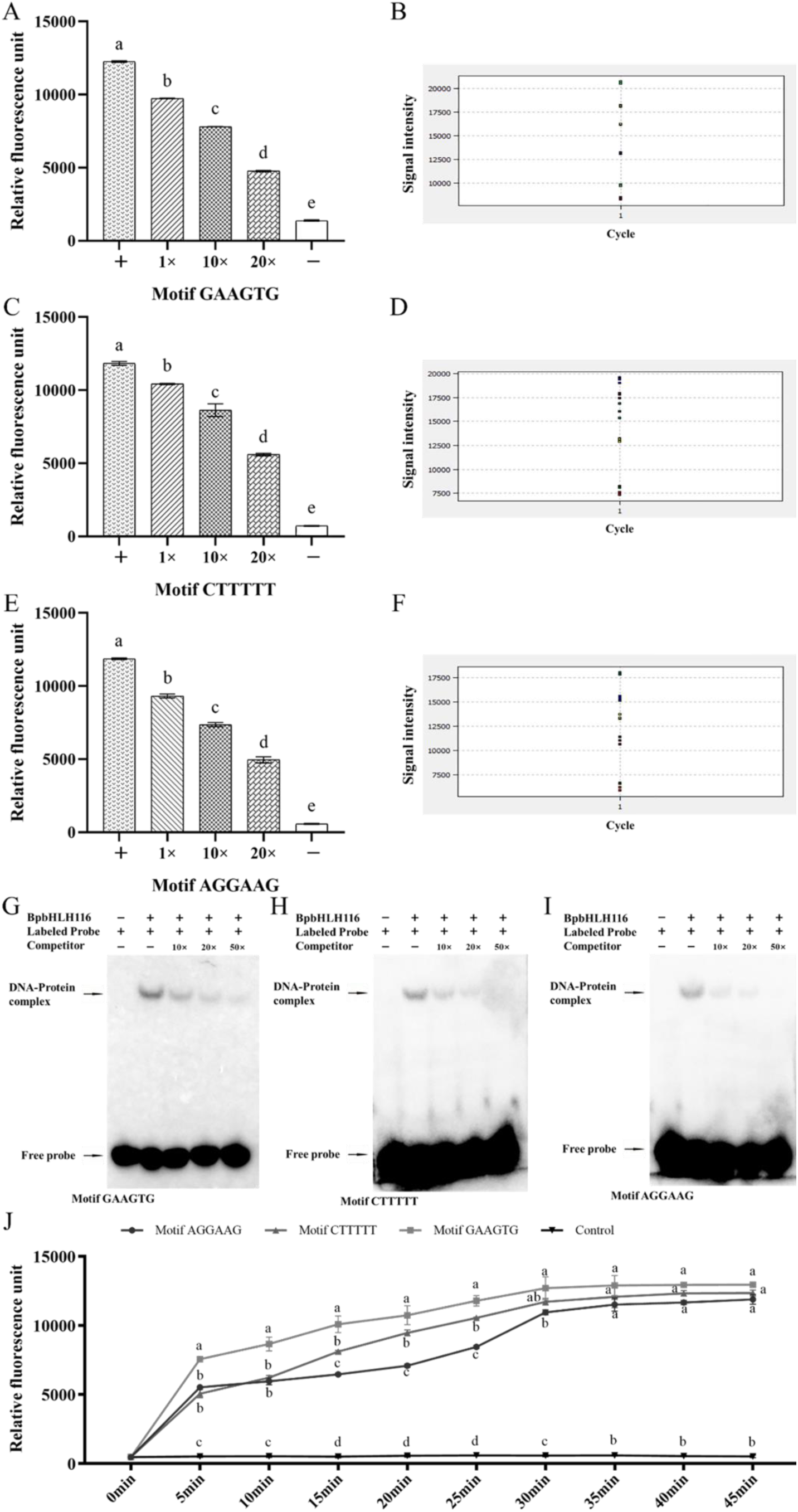
Kinetic Analysis of DNA-Protein Binding Using RP-DBA. A: The binding of the target protein to the DNA motif investigated by RP-DBA. B: the original fluorescence signal intensity graph for the DNA-protein binding, generated by the quantitative PCR instrument. The signal intensity values in panel A are derived from the signal values in panel B after subtracting the background values (using an equal volume of water as a control). C: Confirmation of the binding of the protein to the target DNA motif through Electrophoretic Mobility Shift Assay (EMSA). D: A kinetic analysis of the protein-DNA binding, with binding detection conducted every 5 minutes over a period of 0 to 45 minutes. The statistical significance of intergroup differences was evaluated using one-way analysis of variance (ANOVA) test. The error bars indicate the standard deviations of the mean measurements (n= 3). Letters a-e indicate significant differences (P < 0.01).

We then utilized the RP-DBA method to study the binding kinetics of the bHLH protein with the three DNA elements, establishing ten time points within a 0-45 minute range (with 5-minute intervals) (Figure 3J). The experimental results indicated that the binding levels of all DNA elements increased progressively over time; however, distinct kinetic characteristics were observed among the different motifs. During the initial binding phase (5-10 minutes), bHLH exhibited the fastest binding rate to GAAGTG (from 7555 to 8653, an increase of 14.6%), significantly surpassing its binding to AGGAAG (from 5516 to 5957, an increase of 8.0%) and CTTTTT (from 5062 to 6213, an increase of 22.7%). This suggests that bHLH may have a superior initial recognition efficiency for GAAGTG. Although the initial binding amount of bHLH to CTTTTT was the lowest at this stage, it demonstrated the largest relative increase (22.7%) (Figure 3J).

As the reaction progressed to the mid-phase (20-30 minutes), the binding kinetics began to diverge: the binding of bHLH to GAAGTG slowed down (from 10735 to 12688, Δ4953), while the binding to AGGAAG accelerated (from 7079 to 10947, Δ3868), particularly showing a significant increase during the 25-30 minute interval (an increase of 29.7%). In contrast, the binding of bHLH to CTTTTT exhibited a steady increase (from 9459 to 11712, Δ2252) (Figure 3J).

In the later stages of the reaction (35-45 minutes), bHLH reached binding equilibrium with GAAGTG first (from 12887 to 12941, an increase of only 0.4%), while the binding to AGGAAG (from 11510 to 11884, an increase of 3.2%) and CTTTTT (from 12066 to 12343, an increase of 2.3%) continued to show slight increases, indicating that different elements exhibit varying degrees of binding saturation. Ultimately, the final binding levels exhibited a hierarchical relationship of GAAGTG (12941) > CTTTTT (12343) > AGGAAG (11884) (with no significant differences), indicating that variations in DNA sequences significantly influence the rate of protein-DNA binding (Figure 3J).

### Sensitivity Study of RP-DBA Technology

To investigate the sensitivity of the RP-DBA technology, we compared it with EMSA technique. RP-DBA was performed using a signal amplification arm with a length of 30 bp. In both experiments, we utilized the same amount of protein, which was serially diluted to concentrations of 1, 0.5, 0.2, 0.1, and 0.05 times, and subsequently assessed their binding to probes followed by signal detection. The results indicated that the RP-DBA technology was able to detect a signal significantly above the negative control even at a protein dilution of 0.05 (Figure 4A-B), whereas EMSA reached its minimum detection limit at a dilution of 0.2. This demonstrates that RP-DBA is at least four times more sensitive than EMSA when utilizing an amplification arm that is 30 bp in length(Figure 4C).

**Figure 4:**
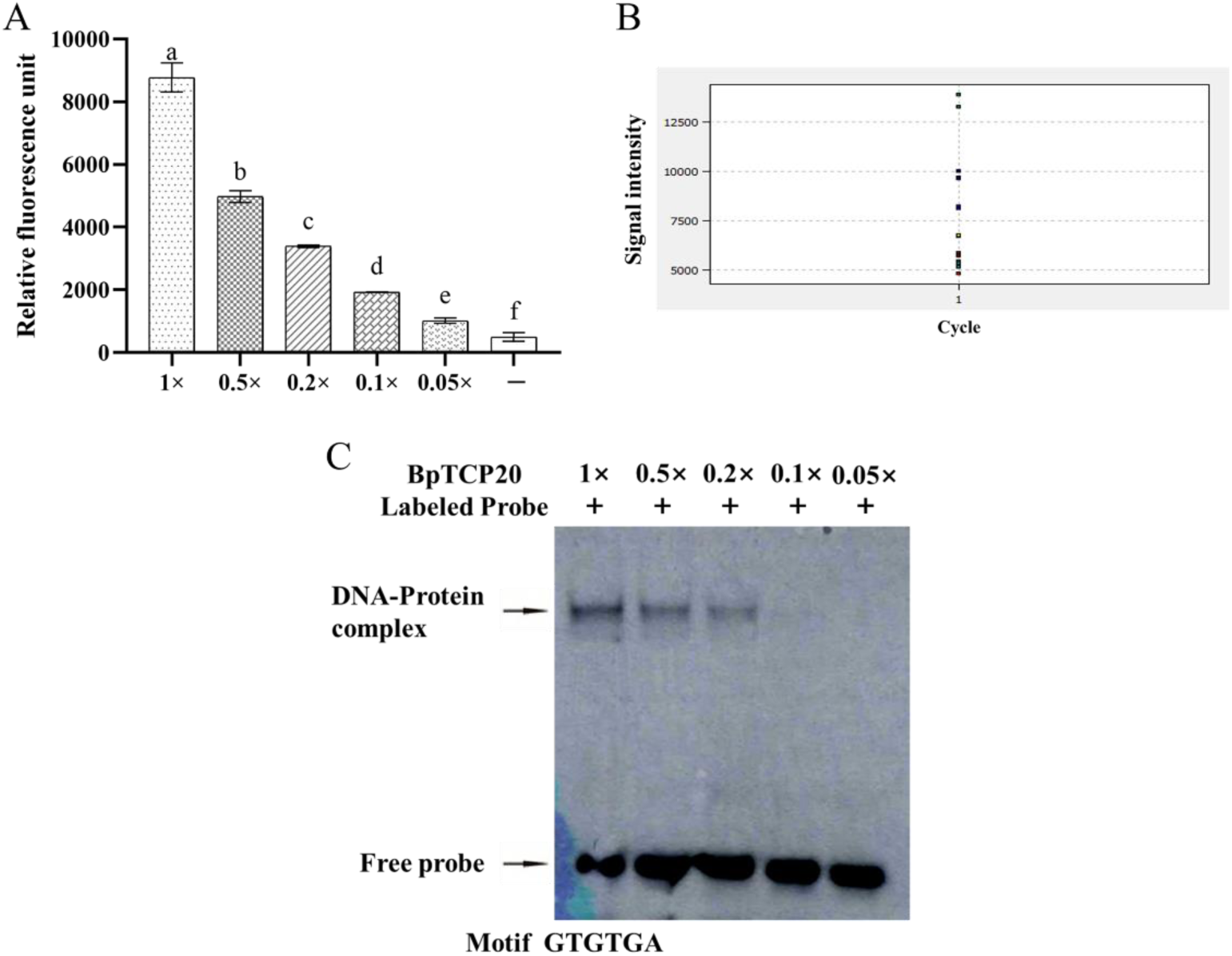
Comparison of Sensitivity Between RP-DBA and EMSA. A and B: The binding of proteins at different concentrations to an excess of DNA probes (A) and the corresponding original fluorescence signal intensity graph for DNA-protein binding, generated by the quantitative PCR instrument (B). The signal intensity values in panel A are obtained by subtracting the background values (using an equal volume of water as a control) from the signal values in panel B. Proteins were diluted in the following ratios: 1x (undiluted), 0.5x, 0.25x, 0.1x, and 0.05x, before conducting the DNA binding experiments. The statistical significance of intergroup differences was evaluated using one-way analysis of variance (ANOVA) test. (C): The EMSA analysis of protein binding to excess DNA probes at different concentrations. The same amount of protein used in RP-DBA was diluted for EMSA, allowing for a comparison of sensitivity between the two methods. The EMSA probes are end-labeled with biotin, and chemiluminescence is utilized to detect the binding of proteins to DNA. The error bars indicate the standard deviations of the mean measurements (n =3). Letters a-e indicate significant differences (P < 0.01).

### Effects of Different Lengths of Single-Stranded DNA Arms on the Sensitivity of RP-DBA

To enhance the signal strength and sensitivity of transcription factor protein-DNA binding, we compared the differences in signal intensity resulting from the addition of single-stranded DNA arms of varying lengths during the PCR amplification process. We selected four proteins for the experiment, conducting PCR amplification without the addition of single-stranded DNA and with the addition of single-stranded DNA arms measuring 30 bp (the length used in previous experiments), 60, and 90 bp. We then assessed the signal intensity and its reproducibility.

The experimental results demonstrated that the addition of single-stranded DNA significantly increased the signal intensity, and this increase was positively correlated with the length of the DNA arms. The signal intensity with 60 bp arms was found to be 1.6 to 2.0 times greater than that of the 30 bp arms, while the signal intensity with 90 bp arms increased approximately 2.4 to 2.8 times compared to the 30 bp arms. Furthermore, the reproducibility of the signals was consistently high across three independent biological replicates (Figure 5A-H).

**Figure 5:**
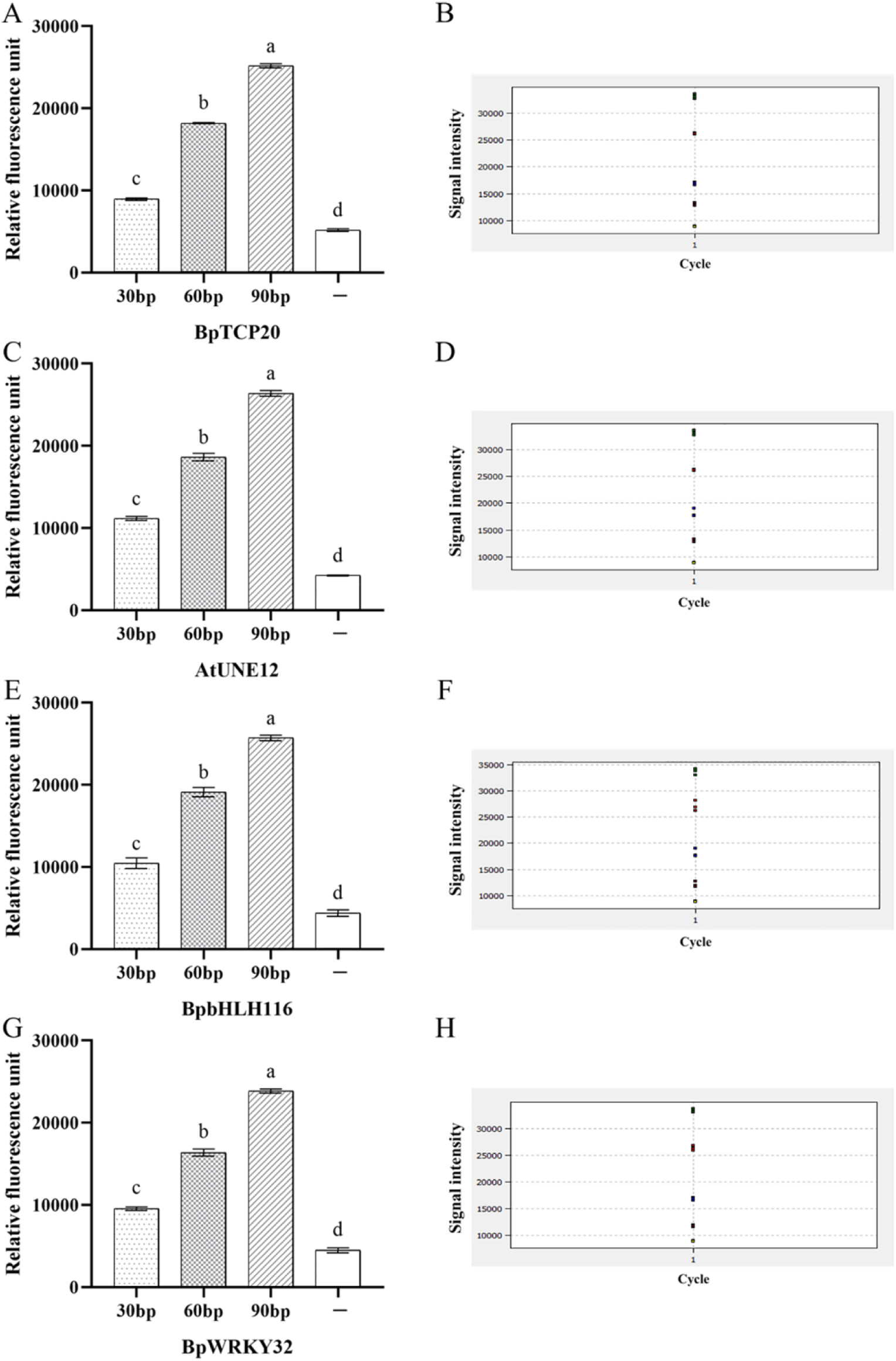
Impact of Signal Amplification Arm Length on Sensitivity in RP-DBA. A, C, E, and G: Four transcription factors, including BpTCP20 (A), AtUNE12 (B), BpbHLH116 (C), and BpWRKY32 (D), which were tested with and without amplification arms of lengths 30 bp, 60 bp, and 90 bp to assess their sensitivity through fluorescence intensity measurements. The signal intensity values in panels A, C, E, and G are calculated by subtracting the background values (using an equal volume of water as a control) from the signal values in panels B, D, F, and H. B, D, F, and H: The original fluorescence signal intensity graphs for DNA-protein binding, generated by the quantitative PCR instrument. The statistical significance of intergroup differences was evaluated using one-way analysis of variance (ANOVA) test.The error bars indicate the standard deviations of the mean measurements (n =3). Letters a-e denote significant differences (P < 0.01).

These findings indicate that the incorporation of single-stranded DNA not only significantly enhances the sensitivity of detection but also improves the reproducibility of experimental results. In summary, the extension of single-stranded DNA arms plays a crucial role in improving the detection efficacy of transcription factor protein-DNA interactions, and increasing the length of these arms can significantly enhance the sensitivity of the RP-DBA method.

## Discussion

This study systematically investigated the specific binding characteristics between transcription factors and DNA motifs based on the RP-DBA technology framework. Experimental data demonstrated that RP-DBA exhibits significant specificity and sensitivity in detecting protein-DNA interactions, with quantitative results showing a high degree of consistency with traditional EMSA (Figure 2A-I, 3A-I). These findings validate the reliability of this method in molecular interaction research.Many techniques are available for the detection and characterization of protein-nucleic acid complexes (Hellman and Fried, 2007). Among these, in vitro analysis methods for protein-DNA interactions primarily include three categories: filter binding and blotting techniques, such as nitrocellulose filter-binding assays (Woodbury et al, 1983) and Southwestern blotting (Bowen et al, 1980); Electrophoretic Mobility Shift Assay (EMSA) techniques (Cozzolino et al, 2021); and structural protection methods, including methylation interference and DNase I footprinting (Gustafsdottir et al, 2007; Tullius et al, 1987). However, these classical methods face technical limitations: the non-specific adsorption rate of nitrocellulose membranes can be as high as 15-30%, leading to false positives; Southwestern blotting involves time-consuming steps (over 8 hours), with a sensitivity limited to the micromolar range; structural protection methods have a success rate of less than 40% for detecting weak binding interactions and require precise control of reaction conditions; although EMSA is widely used, it necessitates complex procedures such as electrophoresis, membrane transfer, and color development, often resulting in experimental cycles exceeding 12 hours.

Building upon the aforementioned three categories of protein-DNA binding methods, we have introduced an additional approach for studying protein-DNA interactions, namely the binding detection method. The RP-DBA technology achieves highly sensitive quantitative detection of protein-DNA interactions by innovatively integrating magnetic bead separation with a qPCR signal amplification system. Its core breakthroughs are primarily reflected in the following aspects: (1) Label-free Universal Detection: RP-DBA employs a sticky-end complementary extension design, converting DNA bound by proteins into qPCR amplification templates. This eliminates the need for traditional probe labeling (such as radioactive or fluorescent labels), making the detection method compatible with standard qPCR instruments and general chemical imaging systems. (2) Rapid Process Driven by Magnetic Bead Separation: Utilizing the “binding-elution” method with Strep-Tactin magnetic beads, RP-DBA reduces the operational time from the traditional 4-6 hours to just 1.5 hours, significantly enhancing the efficiency of high-throughput screening. (3) Dynamic Binding Analysis Capability: RP-DBA utilizes SYBR Green for precise fluorescence quantification of DNA, which markedly improves the sensitivity and accuracy of detection compared to traditional semi-quantitative methods, enabling a deeper analysis of protein-DNA binding kinetics.

The research results indicate that when transcription factors specifically bind to particular DNA sequences, the fluorescence signal gradually diminishes with the addition of competitive probes (Figure 2A-I, 3A-I). This phenomenon effectively distinguishes between specific and non-specific binding. The introduction of competitive probes significantly reduced the fluorescence signal, yet it remained higher than that of the negative control group, further validating the effectiveness and accuracy of RP-DBA in identifying protein-DNA interactions. Additionally, three independent biological replicates demonstrated a high level of consistency (Figures 2-5), further confirming the reliability of this technique.

The RP-DBA technology also enables the kinetic analysis of protein-DNA binding (Figure 3J). Traditional methods for studying protein-DNA interactions, such as Southwestern blotting, nitrocellulose filter-binding assays, and EMSA, often struggle to quantitatively analyze binding affinity and its kinetic characteristics. RP-DBA effectively fills this gap by quantifying the binding capabilities of different proteins to DNA (Figures 2A-I, 3A-I) and revealing significant differences in binding affinity between the same protein and different DNA sequences (Figure 3J). This technique clearly reflects the characteristics of binding kinetics, such as the initial binding rate, mid-to-late stage binding capacity, and final binding saturation (Figure 3J), providing new experimental means for a deeper understanding of the interaction mechanisms between transcription factors and DNA sequences.

The RP-DBA technology exhibits significantly greater sensitivity compared to EMSA. Under identical protein amounts and saturation probe conditions, the fluorescence signal of RP-DBA at 0.05 times the normal protein concentration is markedly higher than that of the negative control (Figures 4A and 4B), while EMSA can barely detect a signal at 0.2 times the concentration (Figure 4C). This indicates that the sensitivity of RP-DBA is at least four times greater than that of EMSA. This analysis was conducted using an amplification arm with a length of 30 bp. However, when a 90 bp amplification arm is used, the sensitivity of RP-DBA can increase by four times (Figure 5A-H), which means that the sensitivity of RP-DBA could reach up to ten times greater than that of EMSA.These results demonstrate that RP-DBA can effectively detect low-abundance protein-DNA interactions that may not be identifiable using EMSA. By converting binding events into quantifiable fluorescence signals, RP-DBA enables the confirmation of weak interactions through statistical analysis, making it particularly well-suited for the study of weakly binding protein-DNA interactions.

RP-DBA further enhances sensitivity by utilizing signal extension with single-stranded DNA that is complementary to the sticky ends of the target probe (Figure 5A-H). In this study, the length of the signal amplification arm was set at 30 bp, and when the length was increased to 60 bp, the signal intensity rose to 1.6-2.0 times that of the 30 bp arm; when the length was further extended to 90 bp, the signal intensity reached 2.4-2.8 times that of the 30 bp arm (Figure 5A-H). Therefore, extending the length of the signal amplification arm can significantly enhance the sensitivity of RP-DBA, a feat that is challenging to achieve with EMSA and other methods. Generally, a 30 bp signal amplification arm length is sufficient to meet experimental needs; however, in specific cases, further sensitivity enhancement can still be achieved by increasing the length of the signal amplification arm.

Although RP-DBA demonstrates simplicity, high specificity, and sensitivity in this study, several key factors must be considered during the experimental process. First, RP-DBA relies on magnetic bead purification of protein-DNA complexes, which carries the risk of nonspecific adsorption. Therefore, it is essential to thoroughly wash the beads during the washing process to minimize background noise, and to strictly adhere to the protocols provided in this experiment. Second, the quantitative analysis of fluorescence signals must take into account the interference of background signals. Studies have shown that even ultrapure water can generate background fluorescence signals in quantitative PCR instruments; thus, it is necessary to design ultrapure water controls and measure their background values to subtract these signals during detection. For weak DNA-protein binding, it is advisable to design 3-5 negative controls to calculate the mean and standard deviation, and to use statistical software to analyze whether there are significant differences between the signal values of DNA-protein binding and the background signal values. This is crucial for accurately detecting weak DNA-protein interactions.

Furthermore, the RP-DBA technique has demonstrated good reproducibility and reliability in preliminary experiments, allowing for the possibility of omitting biological replicates in cases where significant gradient differences in signal formation are observed.

## Conclusion

RP-DBA is a sensitive and efficient method for studying protein-DNA interactions, surpassing EMSA in sensitivity to detect weak binding events at low concentrations. By translating interactions into quantifiable SYBR Green fluorescence via qPCR amplification, it enables kinetic binding analysis and affinity quantification. The integration of Strep-Tactin magnetic bead purification ensures specificity, while its high-throughput capacity supports large-scale screening in functional genomics. This approach addresses critical limitations of conventional methods in resolving transient or low-affinity interactions.

## Supplementary data

### Supplementary Figures

Supplementary Figure 1: Identification of various DNA sequences bound by bHLH through ChIP-seq.

### Supplementary Tables

Table S1: Primers for prokaryotic expression vectors of 4 proteins used in the Rapid Protein-DNA Binding Assay.

Table S2: All probe sequences used for RP-DBA analysis.

Table S3: All probe sequences used for EMSA analysis.

## Data availability

ChIP-Seq data: NCBI SRA PRJNA1247694 (https://www.ncbi.nlm.nih.gov/sra/PRJNA1247694)

